# rawDiag - an R package supporting rational LC-MS method optimization for bottom-up proteomics

**DOI:** 10.1101/304485

**Authors:** Christian Trachsel, Christian Panse, Tobias Kockmann, Witold E. Wolski, Jonas Grossmann, Ralph Schlapbach

**Affiliations:** Functional Genomics Center Zurich Swiss Federal Institute of Technology Zurich I University of Zurich Winterthurerstr. 190, CH-8057 Zurich, Switzerland

**Keywords:** mass spectrometry, visualization

## Abstract

Optimizing methods for liquid chromatography coupled to mass spectrometry (LC-MS) is a non-trivial task. Here we present `rawDiag`, a software tool supporting rational method optimization by providing MS operator-tailored diagnostic plots of scan level metadata. `rawDiag` is implemented as R package and can be executed on the command line, or through a graphical user interface (GUI) for less experienced users. The code runs platform independent and can process a hundred raw files in less than three minutes on current consumer hardware as we show by our benchmark. In order to demonstrate the functionality of our package, we included a real-world example taken from our daily core facility business.

## 1 Introduction

Over the last decade, liquid chromatography coupled to mass spectrometry (LC-MS) has evolved into the method of choice in the field of proteomics.^1–5^ During a typical bottom up LC-MS measurement, a complex mixture of analytes is separated by a liquid chromatography system that is connected to a mass spectrometer (MS) through an ion source interface. The analytes which elute from the chromatography system over time are converted into a beam of ions in this interface and the MS records from this ion beam a series of mass spectra containing detailed information on the analyzed sample.^6,7^ These mass spectra, as well as their metadata, are considered as the raw measurement data and usually recorded in a vendor specific binary format. During a measurement, the mass spectrometer applies internal heuristics which enables the instrument to adapt to sample properties like sample complexity or amount in near real time. Still, method parameters controlling these heuristics, need to be set prior to the measurement. For an optimal measurement result, a carefully balanced set of parameters is required, but their complex interactions with each other make LC-MS method optimization a challenging task.

Here we present `rawDiag`, a platform independent software tool implemented in R that supports LC-MS operators during the process of empirical method optimization. Our work builds on the ideas of the discontinued software “rawMeat” (vastScientific). Our application is currently tailored towards spectral data acquired on Thermo Fisher Scientific instruments (raw format), with a special focus on Orbitrap mass analysers (Exactive or Fusion instruments). These instruments are heavily used in the field of bottom-up proteomics in order to analyse complex peptide mixtures derived from enzymatic digests of proteomes. `rawDiag` is meant to run post mass spectrometry acquisition, optimally as interactive R shiny application and produces a series of diagnostic plots visualizing the impact of method parameter choices on the acquired data across injections. If static reports are required, pdf files can be generated using R markdown. The visualizations generated by `rawDiag` can be used in an iterative method optimization process (see Figure 1) where an initial method is tested, analyzed and based on these results a hypothesis can be formulated to optimize the method parameters. The same sample is then re-analyzed with the optimized method and the data can enter the refinement loop again until the operator is satisfied with the found set of method parameters for his type of sample.

**Figure 1:**
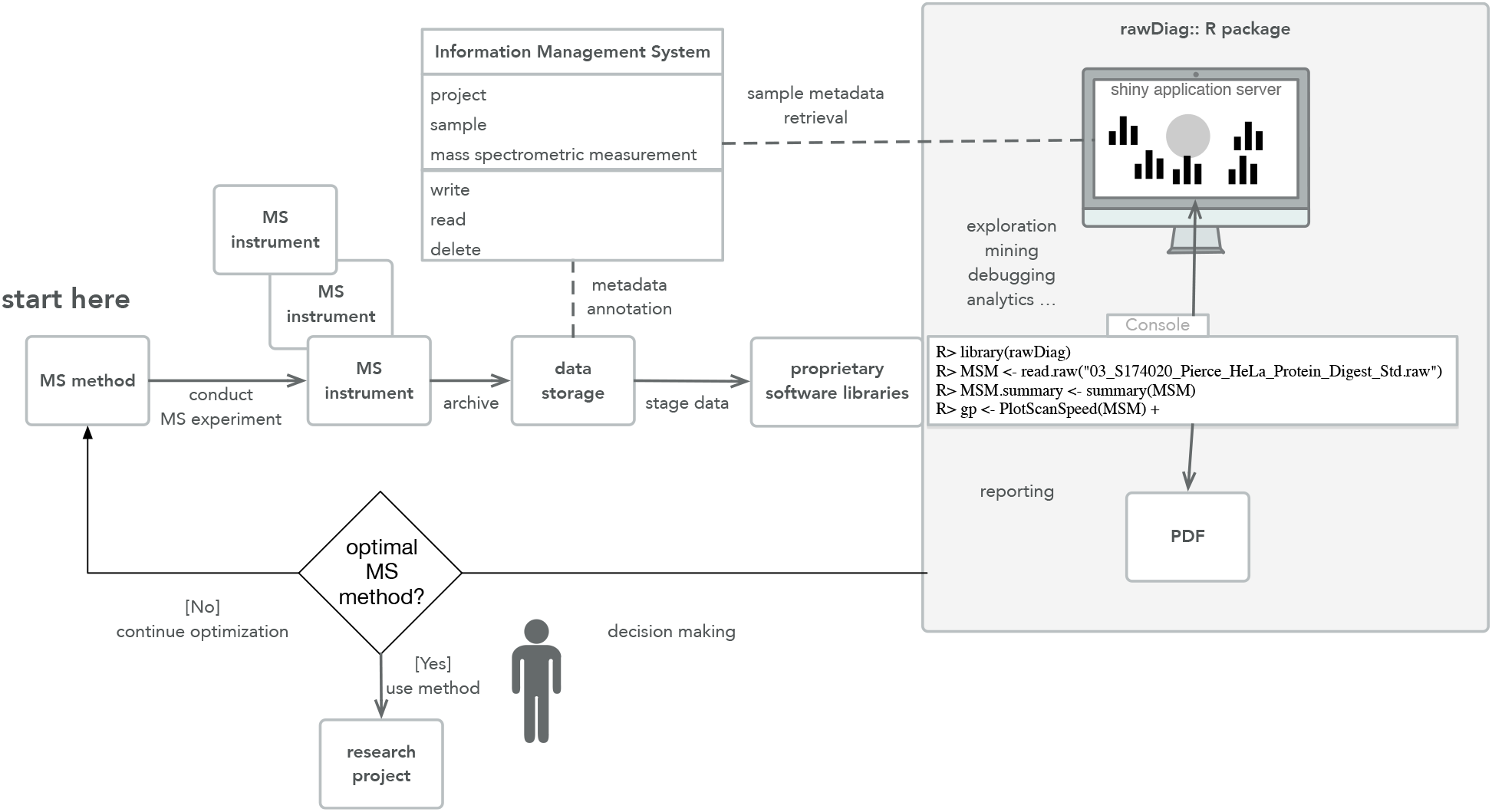
The schema displays the f eedback loop of the optimization process of an LC-MS method using `rawDiag` (see the grey box on the right). The optimization starts (“start here”) with an initial method. Scan data is recorded using the initial method and the information stored as raw instrument data. `rawDiag` reads the scan metadata data and visualizes the method characteristics. Based on this analysis, the mass spectrometer operator can optimize the instrument method. Optionally, `rawDiag` could also operate on top of a lab information management system(LIMS).

In this manuscript we present the architecture and implementation of our tool. We provide example plots, show how plots can redesigned to meet different requirements and discuss the application based on a use cases.

## 2 Experimental Procedures

### 2.1 Architecture

`rawDiag` acts as an interface to vendor specific software libraries that are able to access the spectrum-level metadata contained in a mass spectrometer measurement. The package provides support for R command line, interactivity through R shiny and pdf report generation using R markdown. A rough overview of the architecture can be seen in Figure 1.

### 2.2 Implementation

The entire software is implemented as R package providing a full documentation and includes example data. All diagnostic plots are generated by R functions using the ggplot2^8^ graphical system, based on “The Grammar of Graphics”.^9^ The package ships with an adapter function *`read.raw`* which returns an R *`data.frame`* object from the raw data input file. In its current implementation, the adapter functions default input method is set for reading Thermo Fisher Scientific raw files, using a `C#` programmed executable^1^, based on the platform-independent RawFileReader.Net assembly^2^. Since in general more than one mass spectrometry file is loaded and visualized, the adapter function supports multiprocessor infrastructure through the parallel R package. In order to be flexible with the entire variety of instruments, we implemented the two utility functions *`is. rawDiag`* and *`as. rawDiag`*. While the *is. rawDiag* function checks if the input object fulfills the requirements of the package’s diagnostic plot functions, the *`as.rawDiag`* method coerce the object into the right format by deriving missing values if possible, otherwise filling missing columns with NA values.

#### 2.3 Visualization

This package is providing several plot functions tailored towards mass spectrometry data. A list of the implemented plot functions with a short description can be found in Table 1. An inherent problem of visualizing data is the fact that depending on the data at hand certain visualizations lose their usefulness (e.g. overplotting in scatter plot if too many data points are present). To address this problematic, we implemented most of the plot functions in different versions inspired by the work of Cleveland,^10^ Sarkar^11^ and Wickham.^8^ The data can be displayed in trellis plot manner using the faceting functionality of ggplot2 (see Figure 2A). Alternatively, overplotting using color coding (Figure 2B) or violin plots based on descriptive statistics values (Figure 2C) can be chosen. This allows the user to interactively change the appearance of the plots based on the situation at hand. E.g. a large number of files are best visualized by violin plots giving the user an idea about the distribution of the data points. Based on this a smaller subset of files can be selected and visualized with another technique.

**Table 1:**
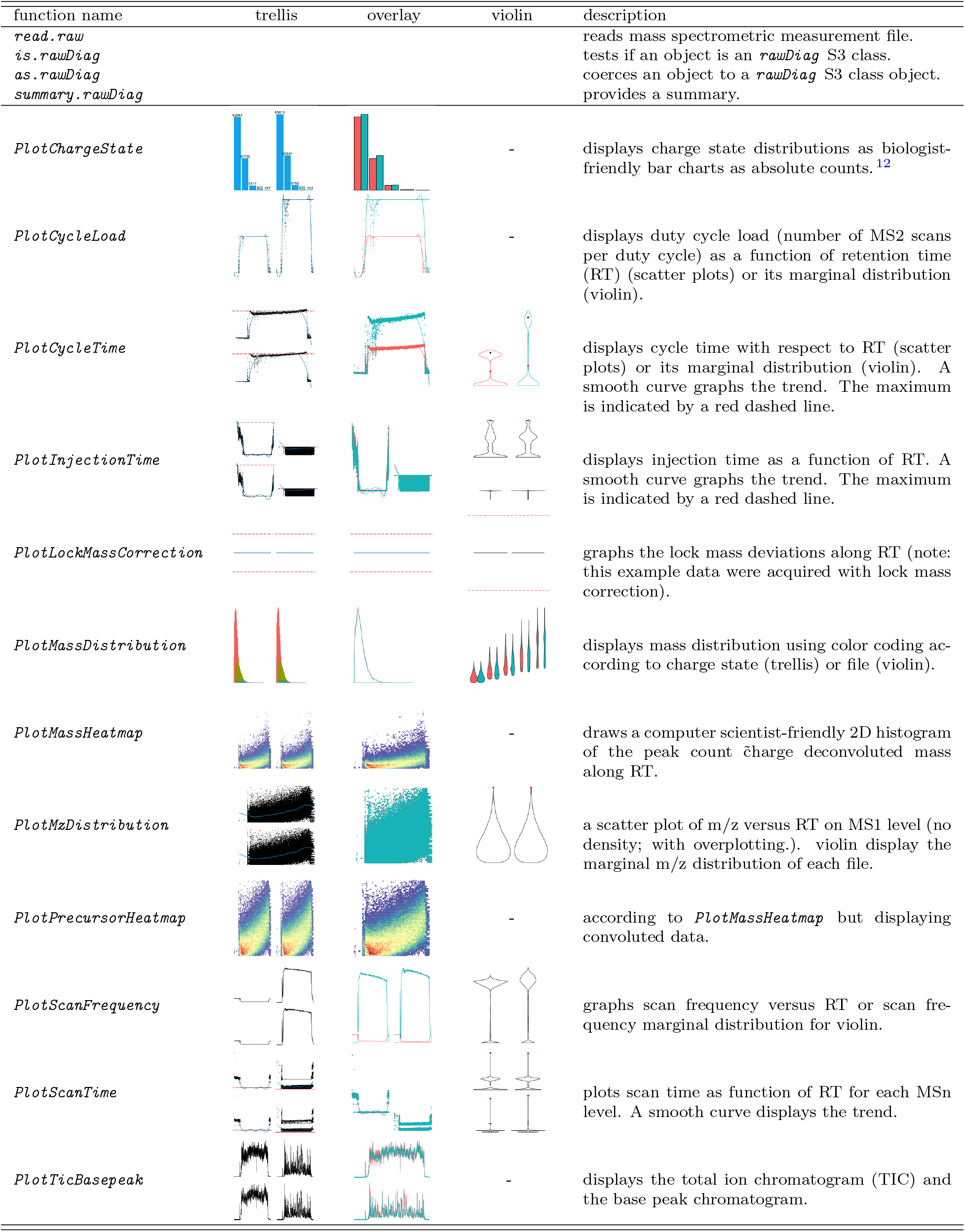
The `rawDiag` cheatsheet lists the functions of the package using a subset of the provided ‘WU163763’ dataset. Each thumbnail gives an impression of the plot function’s result. The column names ‘trellis’, ‘overlay’ and ‘violin’ were given as method attribute. Marginal distribution plots for discrete response variables are not supported.

| function name | trellis | overlay | violin | description |
| --- | --- | --- | --- | --- |
| `read.raw` |  |  |  | reads mass spectrometric measurement file. |
| `is.rawDiag` |  |  |  | tests if an object is an `rawDiag` S3 class. |
| `as.rawDiag` |  |  |  | coerces an object to a `rawDiag` S3 class object. |
| `summary.rawDiag` |  |  |  | provides a summary. |
| `PlotChargeState` |  |  | - | displays charge state distributions as biologist-friendly bar charts as absolute counts. <sup>12</sup> |
| `PlotCycleLoad` |  |  | - | displays duty cycle load (number of MS2 scans per duty cycle) as a function of retention time (RT) (scatter plots) or its marginal distribution (violin). |
| `PlotCycleTime` |  |  |  | displays cycle time with respect to RT (scatter plots) or its marginal distribution (violin). A smooth curve graphs the trend. The maximum is indicated by a red dashed line. |
| `PlotInjectionTime` |  |  |  | displays injection time as a function of RT. A smooth curve graphs the trend. The maximum is indicated by a red dashed line. |
| `PlotLockMassCorrection` |  |  |  | graphs the lock mass deviations along RT (note: this example data were acquired with lock mass correction). |
| `PlotMassDistribution` |  |  |  | displays mass distribution using color coding according to charge state (trellis) or file (violin). |
| `PlotMassHeatmap` |  |  | - | draws a computer scientist-friendly 2D histogram of the peak count charge deconvoluted mass along RT. |
| `PlotMzDistribution` |  |  |  | a scatter plot of m/z versus RT on MS1 level (no density; with overplotting). violin display the marginal m/z distribution of each file. |
| `PlotPrecursorHeatmap` |  |  | - | according to `PlotMassHeatmap` but displaying convoluted data. |
| `PlotScanFrequency` |  |  |  | graphs scan frequency versus RT or scan frequency marginal distribution for violin. |
| `PlotScanTime` |  |  |  | plots scan time as function of RT for each MSn level. A smooth curve displays the trend. |
| `PlotTicBasepeak` |  |  | - | displays the total ion chromatogram (TIC) and the base peak chromatogram. |

**Figure 2:**
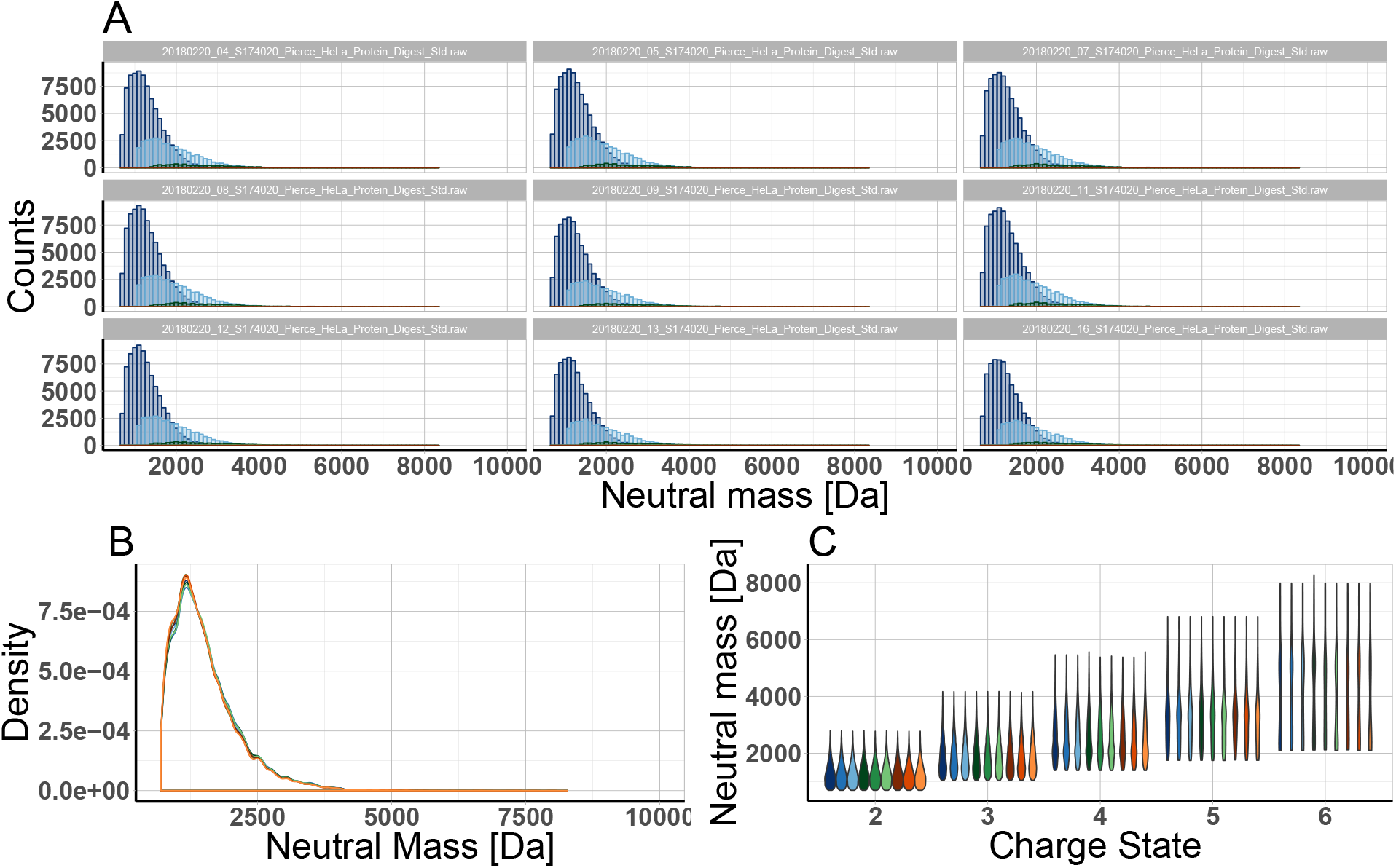
Concurrent metadata visualization applying *`PlotMassDistribution`* to nine raw files acquired in DDA mode (sample was 1*μ*g HeLa digest.) **A)** method trellis **B)** method overlay **C)** method violin.

To benefit from the grammar of graphics, e.g., adapt y-axis scaling, change axis labels, add title or subtitles, each of the implemented plot functions always returns the *`ggplot`* object. Due to the implementation of this design pattern those *`ggplot`* objects can be further altered by adding new layers allowing a customization of the plots if needed. The following R code snippet produces the three plots shown in Figure 2 and demonstrates the described feature of modifying an existing *`ggplot`* object by eliminating the legend in the last two plots.

~~~
R> library(rawDiag)
~~~

~~~
R> WU163763 <- getWU163763()
~~~

~~~
R> PlotMassDistribution(WU163763)
~~~

~~~
R> PlotMassDistribution(WU163763, method = ‘overlay’) +
~~~

~~~
+ theme(legend.position = ‘none’)
~~~

~~~
R> PlotMassDistribution(WU163763, method = ‘violin’) +
~~~

~~~
+ theme(legend.position = ‘none’)
~~~

The interactivity of the visualizations is achieved by an implementation of the plot functions into an R shiny applications. Static versions of the plots can be easily generated by the provided R markdown file that allows the generation of pdf reports.

#### 2.4 Evaluation

We tested the performance of our approach by running an scan information throughput benchmark as a function of the number of used processes on a Linux server and an Apple MacBook Pro. The hardware specifications are listed in Table 2.

**Table 2:** Summary of the hardware specifications.

| specs | Linux Server | Apple MacBook Pro 2017 |
| --- | --- | --- |
| number of cores | 64 | 8 |
| CPU | Intel(R) Xeon(R) CPU E5-2698 v3 @ 2.30GHz | 2.9GHz Intel Core i7 |
| disk | RAID Module RMS25CB080 | SSD SM1024L |
| filesystem | XFS | APFS |
| OS | SMP Debian 3.16.43-2+deb8u2 | Darwin Kernel Version 17.4.0 |
| Mono JIT compiler version | 5.8.0.127 | 5.2.0.224 |

As benchmark data, we downloaded the raw files described in^13^ on the filesystem. For the benchmark we limited the input to 128 files, corresponding to two times the available number of processor cores of the Linux system. The data has an overall file size of 95 GBytes and contains 4×149’326 individual mass spectra in total.

The left plot in Figure 3 depicts the overall runtime in dependency of the number of used CPUs for five repetitions starting with 64 cores to avoid caching issues. The right scatter plot in Figure 3 is derived from the overall runtime and illustrates the scan information throughput in dependency of the number of used process cores.

**Figure 3:**
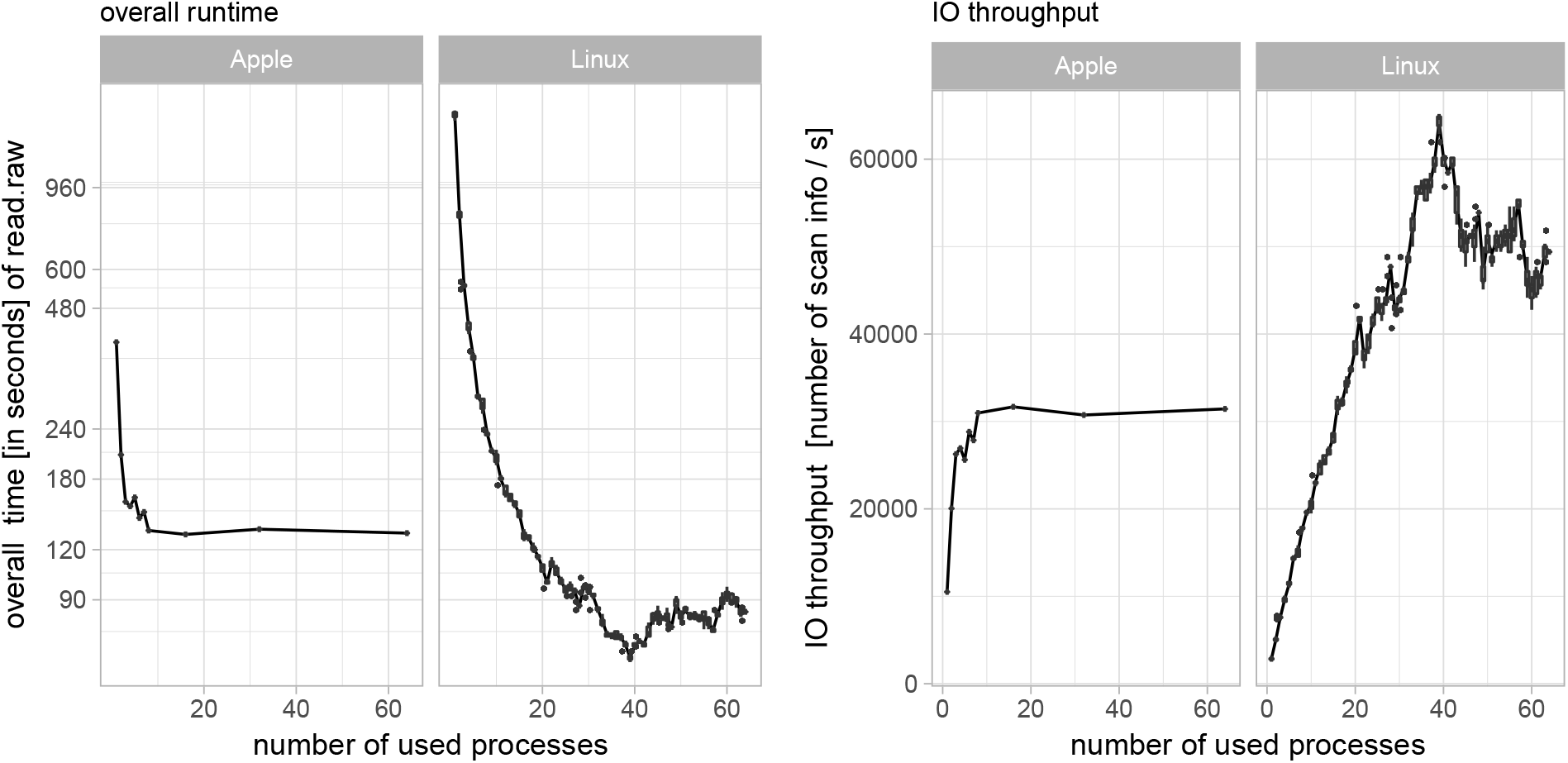
Benchmark - The left plot shows the overall logarithmic scaled runtime of 128 raw files. The graphic on the right side shows the thereof derived IO throughput as scan information per second. The plots illustrate that both systems, server, and laptop, can analyze 95GB of instrument data within less than three minutes.

The best performance on our system is achieved by using 39 CPUs having an performance of reading 64×833 scan information per second.

### 3 Results and discussion

Our application `rawDiag` acts as an interface to file reader libraries from mass spectrometry vendors. These libraries are able to access the scan data as well as the scan metadata stored in the proprietary file formats. In its current configuration, `rawDiag` is able to read data from Thermo Fischer Scientific raw files via a `C#` executable. This executable is extracting the information stored in the raw file via the platform-independent RawFileReader.Net assembly. To avoid writing to the disk, the information is directly fed into an R session using the *`pipe`* command. The data integrity is checked by the *`is.rawDiag`* function and coerced by the *`as.rawDiag`* function into the proper format for the plot functions if required. As soon as the data is extracted and loaded into the R session, the different plot functions can be called upon the data for the visualizations of LC-MS run characteristics. In the envisioned method optimization pipeline (see Figure 1) a test sample which mimics the actual research sample as close as possible is analyzed with an initial method. After the analysis is finished, the acquired data can be visualized by our application. Based on the visualized run characteristics a hypothesis for the method optimizations can be formulated and the optimized methods can again be used to analyze the test sample. A use case example of this process will be discussed in the following paragraph. In the interactive mode, the application runs as an R shiny server and generates a summary table of all loaded data allowing to get an overview in a single glance. The user is provided with a series of plots which provide a rational basis for optimizing method parameters during the iterative process of empirical mass spectrometry method optimization. In order to be flexible towards different situations where a single visualization technique might lose its usability, most plot functions can be called in three different versions. This allows to circumvent overplotting issues or helps to detect trends when multiple files are loaded. A list of the currently implemented plot functions can be found in Table 1 and the flexibility of choosing different visualization styles is depicted in Figure 2.

#### 3.1 Use case — Optimize data dependent analysis

Starting from an initial method template based on the work of Kelstrup et al.,^13^ we analyzed 1 *μ*g of a commercial tryptic HeLa digest on a Q-Exactive HF-X instrument using a classical shotgun heuristics. Subsequently, the resulting raw data was mined using `rawDiag`. Inspection of the diagnostic plots suggested that analysis time was not optimally distributed between the different scan levels (precursor and fragment ions). To test this hypothesis, we ramped the parameter controlling the number of dependent scans per instrument cycle (TopN), in two steps and applied the resulting methods by analyzing the same material in technical triplicates. Visualization applying `rawDiag` confirmed that all three methods exploit the max. number of dependent scans (18, 36 and 72) during the separation phase of the gradient (see Figure 4B). Concurrently, the MS2 scan speed increased from ≈30 to ≈36 and ≈40 Hz respectively (see Figure 4A). Using the modified methods, the instrument is spending 5 and 10 min more time on MS2 scans during the main peptide elution phase, comparing the methods to the initial “Top18” method (see Figure 4E). As a concomitant effect the average cycle load (number of MS2 scans per cycle) increased from 10 (“Top18”) to 16 and 21 (“Top36” and “Top72”, respectively). These optimized methods not only showed better run characteristics but ultimately also resulted in more peptide and protein identifications as shown in Figure 4D and 4D. (Data searched by Sequest through ProteinDiscoverer agains a human database applying standard search parameters and filtered for high confidence)

**Figure 4:**
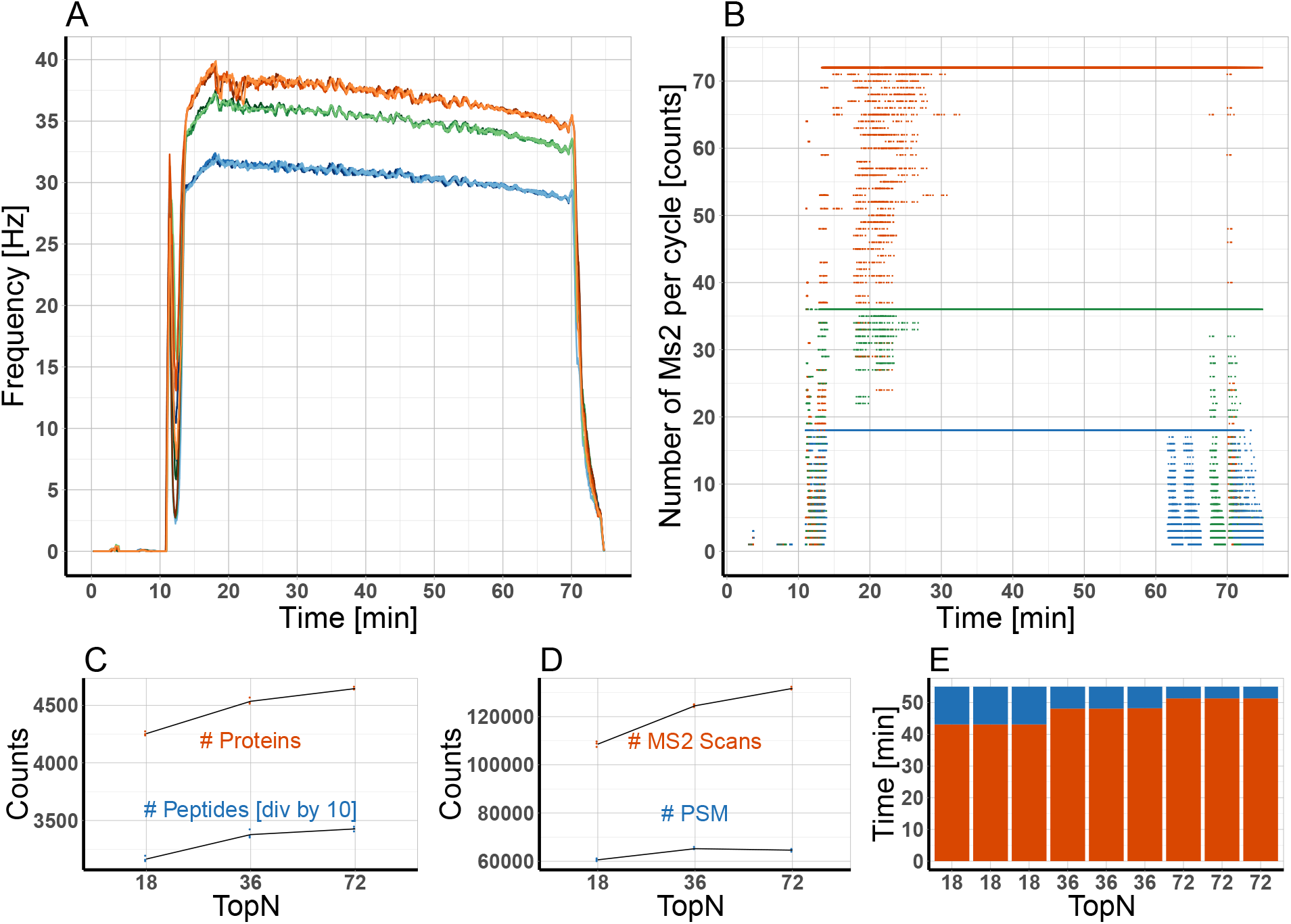
**A)** Moving average of the scan speed of triplicate measurements of “Top18” (blue), “Top36” (green) and “Top72” (orange). **B)** Number of MS2 scans for each scan cycle for “Top18” (blue), “Top36” (green) and “Top72” (orange). **C)** Number of Proteins (orange) and peptides (blue) for the different TopN settings (note: number of peptides is divided by 10 in this plot due to scaling reasons). **D)** Number of PSM (blue) and MS2 scans (orange) for the different TopN settings. **E)** Time spent on MS1 (blue) and MS2 (orange) for the different TopN settings. Time range for calculation is the elution phase of the peptides between 15-70 min.

Interestingly, the number of peptide spectrum matches (PSM) is decreasing in the “Top72” method compared with the “Top36”. Based on this one could formulate a new hypothesis: the “Top72” method is sampling the precursors to such a deep level, that we reach an injection time limit for many low abundant species (MS2 quality is suffering from low amount of ions). To test this, the methods could now be further fine tuned by reducing the number of MS2 from 72 to a lower number but at the same time increase the injection time to increase the spectra quality.

#### 3.2 Related work

To our knowledge the only alternative tool that is able to extract and visualize metadata from raw files on single scan granularity is rawMeat (Vast Scientific). Unfortunately, it was discontinued years ago and built on the meanwhile outdated MSFileReader libraries from Thermo Fisher Scientific (MS Windows only). This implies that it does not fully support the latest generation of qOrbi instruments. Other loosely related tools^14–17^ are tailored towards longitudinal data recording and serve the purpose of quality control (monitoring of instrument performance) rather than method optimization.

### 4 Conclusion

In this manuscript, we presented `rawDiag` an R package to visualize characteristics of LC-MS measurements. Through its diagnostic plots, `rawDiag` supports scientists during empirical method optimization by providing a rational base for choosing appropriate data acquisition parameters. The software is interactive and easy to operate through an R shiny GUI application, even for users without prior R knowledge. More advanced users can fully customize the appearance of the visualizations by executing their own code from the R command line. This also enables `rawDiag` to be customized and implemented into more complex environments, e.g., data analysis pipelines embedded into LIMS systems. In its current implementation, the software is tailored towards the Thermo Fisher Scientific raw file format, but its architecture allows easy adaptation towards other mass spectrometry data formats. An interesting showcase would be the novel Bruker tdf 2.0 format (introduced for the timsTOF Pro), where scan metadata is not “hidden” in proprietary binary files, but stored in an open SQLite database directly accessible to R. In the future, we plan to extend `rawDiag` by allowing users to link additional metadata not originally logged by the instrument software (derived meta-data), but created offline by external tools. A simple, but very useful example, is to scan metadata created by search engines. These typically links scans to similarity scores, peptide assignments, and their corresponding probabilities. This would allow visualizing assignment rates and score distributions across injections. Having the peptide assignments at hand will open the door for chained metadata usage, for instance by linking a scan over the amino acid sequence of the identified peptide to physicochemical properties like hydrophobicity, iRT scores, or MW. Such derived metadata can then be compared to primary metadata like empirical mass or RT. Linking primary and derived metadata will also clear the way to big data applications similar to MassIVE^3^ but bypassing the necessary conversion to open data formats like mzML. This is beneficial since the conversion process does not preserve all useful primary metadata.

### Supporting Information Available

The package vignette as well as the R package itself, a Dockerfile which build the entire architecture from scratch, is accessible through a git repository under the following URL: https://github.com/protViz/rawDiag.

A demo system including all data shown in this manuscript is available through http://fgcz-ms-shiny.uzh.ch:8080/rawDiag-demo.

## Footnotes

1 A Docker recipe for the entire build process of the `C#` based executable also ships with the R package.

2 http://planetorbitrap.com/rawfilereader December 2017

3 https://massive.ucsd.edu/ProteoSAFe/static/massive.jsp, March 2018

## A R Session information

An overview of the package versions used to produce this document are shown below.

- R version 3.4.2 (2017-09-28), `x86_64-apple-darwin15.6.0`
- Locale: `en_US.UTF-8/en_US.UTF-8/en_US.UTF-8/C/en_US.UTF-8/en_US.UTF-8`
- Running under: `macũS High Sierra 10.13.4`
- Matrix products: default
- BLAS: `/Library/Frameworks/R.framework/Versions/3.4/Resources/lib/libRblas.0.dylib`
- LAPACK: `/Library/Frameworks/R.framework/Versions/3.4/Resources/lib/libRlapack.dylib`
- Base packages: base, datasets, graphics, grDevices, methods, stats, utils
- Other packages: bindrcpp 0.2, dplyr 0.7.4, ggplot2 2.2.1, purrr 0.2.4, rawDiag 0.0.2, readr 1.1.1, tibble 1.3.3, tidyr 0.6.3, tidyverse 1.1.1
- Loaded via a namespace (and not attached): assertthat 0.2.0, bindr 0.1, broom 0.4.2, cellranger 1.1.0, colorspace 1.3-2, compiler 3.4.2, forcats 0.2.0, foreign 0.8-69, glue 1.2.0, grid 3.4.2, gtable 0.2.0, haven 1.0.0, hexbin 1.27.1, hms 0.3, httr 1.2.1, jsonlite 1.5, labeling 0.3, lattice 0.20-35, lazyeval 0.2.0, lubridate 1.6.0, magrittr 1.5, mnormt 1.5-5, modelr 0.1.0, munsell 0.4.3, nlme 3.1-131, parallel 3.4.2, pkgconfig 2.0.1, plyr 1.8.4, psych 1.7.3.21, R6 2.2.2, Rcpp 0.12.12, readxl 1.0.0, reshape2 1.4.2, rlang 0.1.2, rvest 0.3.2, scales 0.5.0, stringi 1.1.5, stringr 1.2.0, tools 3.4.2, xml2 1.1.1

